# On Correlation between Structural Properties and Viral Escape Measurements from Deep Mutational Scanning

**DOI:** 10.1101/2022.02.17.480939

**Authors:** Leili Zhang, Giacomo Domeniconi, Chih-Chieh Yang

## Abstract

Encouraged by recent efforts to map responses of SARS-CoV-2 mutations to various antibody treatments with deep mutational scanning, we explored the possibility of tying measurable structural contact information from the binding complexes of antibodies and their targets to experimentally determined viral escape responses. With just a single crystal structure for each binding complex, we find that the average correlation coefficient *R* is surprisingly high at 0.76. Our two methods for calculating contact information use binary contacts measured between all residues of two proteins. By varying the parameters to obtain binary contacts, we find that 3.6 Å and 7 Å are pivotal distances to toggle the binary step function when tallying the contacts for each method. The correlations are improved by short simulations (∼25 ns), which increase average *R* to 0.78. With blind tests using the random forest model, we can further improve average *R* to 0.84. These easy-to-implement measurements can be utilized in computational screening of viral mutations that escape antibody treatments and potentially other protein-protein interaction problems.

## Introduction

Antibody treatment is a powerful, historically proven tool to fight diseases caused by infections using human body’s own armory (*1*). With the advancement of technologies such as mouse hydridoma (*2*), phage display (*3*), and transgenic mouse (*4, 5*), hundreds of successful therapeutic antibodies have been developed to treat various types of cancer (*6*), autoimmune diseases (*7*) and viral infections (*8*). Of particular note in the field is the quick development of new antibodies designed to treat COVID-19 (*9–15*). However, as the pandemic drags on, new SARS-CoV-2 variants have developed resistance to several of these antibodies (*16–19*), calling for a new round of antibody development or antibody improvement. Among the emerging ways to optimize antibodies tailored for specific mutants, Deep Mutational Scanning (DMS) (*20*) has shown promise, predicting hundreds of possible viral escapes given a set of antibody treatments (*21–23*). Benefiting from high-throughput sequencing technology (*24*) and the advancement in data analysis algorithms in recent years (*25*), DMS is used to understand protein properties, protein evolution, engineer proteins, and to solve protein structures (*26, 27*). However, the mass volume of information collected from DMS has been daunting to the developers (*20*) as they try to achieve “simple” outcomes such as antibody optimization. In 2021, Starr et al. showed that, when DMS is used with computational modeling tools such as ROSETTA (*23, 28*), such a task is possible. Their work offers an initial demonstration how DMS and molecular modeling can be combined to accelerate antibody designs with plausible results.

Ideally, if we can replace DMS with a computational method that has comparable accuracy, the antibody design process proposed in Starr et al. can be fully computerized, thus significantly reducing the total cost of the process. Several studies have attempted this since the publication of Starr et al. For example, Pavlova et al. showed that traditional free energy perturbation (FEP) results agree well with DMS (*29*). However, FEP calculations on a single mutation normally requires at least 5 replicas of 20 ns simulations to guarantee accuracy. This has prevented FEP from being used to fast-screen hundreds of mutations which are required by antibody designs. Note that Pavlova et al. (*29*) also presented a machine learning (ML) method that distinguishes the interactions between SARS-CoV receptor binding domain (RBD, Figure 1A) and SARS-CoV-2 RBD when bound with human angiotensin-converting enzyme 2 (ACE2, Figure 1B). However, the labels used to train the model were not related to the antibody designs. Capponi et al. (*30*) directly trained the image-based ML model on the DMS data and achieved over 90% accuracy when predicting whether a mutation strengthens or weakens RBD-ACE2 binding. A drawback of this method is that the prediction of one mutation requires one molecular dynamics (MD) simulation (at least 200 ns), therefore demanding a similar amount of computational resources as FEP calculations. Our group has also applied CASTELO (a contact-matrix-based lead optimization tool) (*31, 32*) to optimize existing antibodies to treat SARS-CoV-2 mutants (*33*). Other studies have attempted to use sequence information to train the ML models against DMS data and yielded plausible results (*34*).

**Figure 1:**
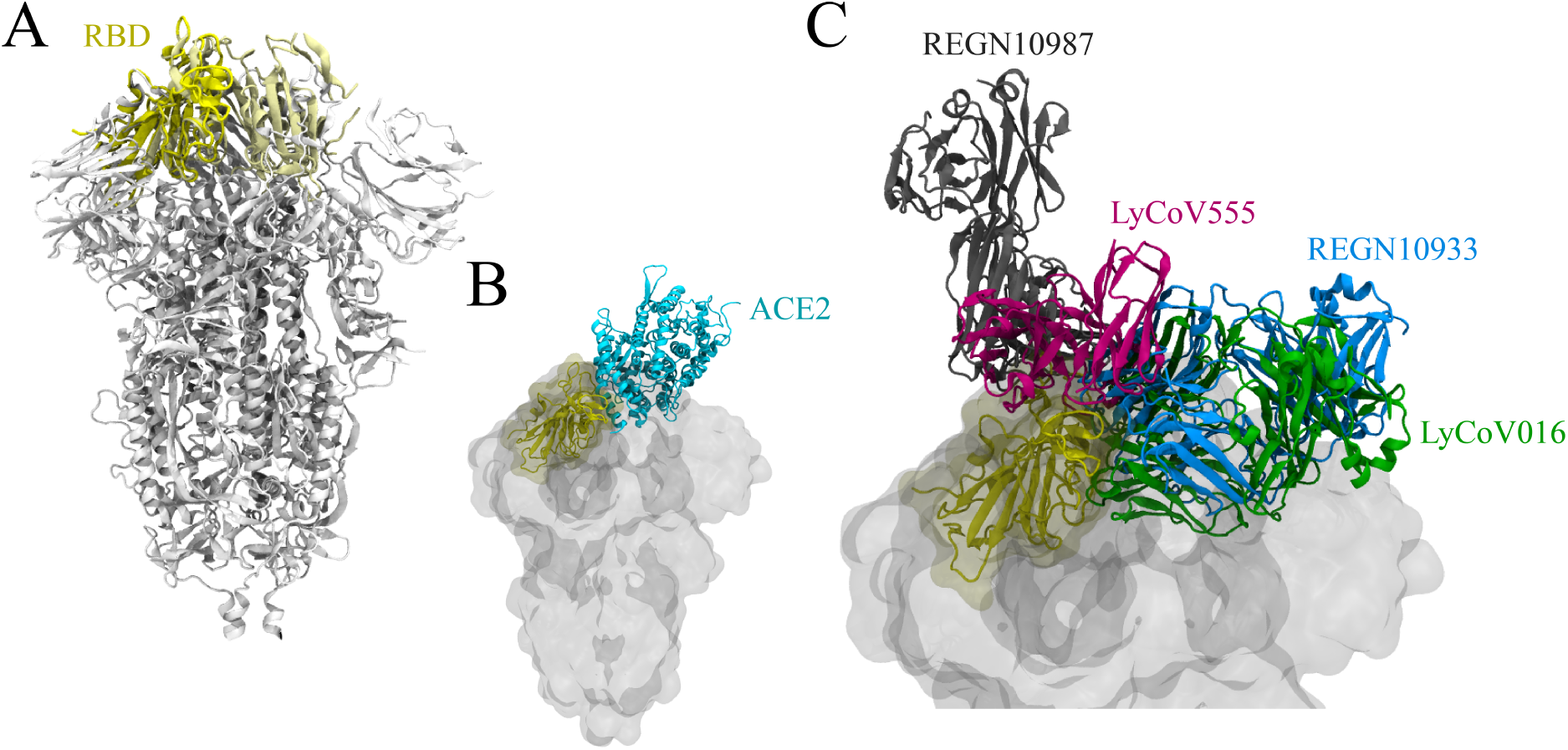
(A) Structure of SARS-CoV-2 spike protein (white), with RBD highlighted in yellow (PDB ID: 6VXX (*48*)); (B) Crystal structure where RBD is found to interact with hACE2 (PDB ID: 6M0J), overlaid with SARS-CoV-2 S protein. (C) Crystal structures where RBD binds with antibodies, overlaid with SARS-CoV-2 spike protein: LyCoV016 in green (PDB ID: 7C01), LyCoV555 in magenta (PDB ID: 7KMG), REGN10933 in blue (PDB ID: 6XDG), and REGN10987 in gray (PDB ID: 6XDG).

Inspired by the successes from structure-based ML methods, we asked “what makes structural-based ML methods work?”. To answer this question, we broke down the steps used in CASTELO (*31,32*) and found that simple contact properties derived from protein complex structures highly correlate with the DMS viral escape measurements from Starr et al. (*21, 22*) for four antibody-RBD complexes (the four antibodies examined in this study are LyCoV016, LyCoV555, REGN10933 and REGN10987, shown in Figure 1C). We were able to show that, with just four crystal binding complexes of the wildtypes, structural properties such as contact percentage and total contacts derived from the frame-wise contact matrices (method summarized in Figure 2) correlate surprisingly well with DMS viral escape measurements (which are derived from hundreds of DMS measurements). Furthermore, the correlations are improved by short molecular dynamics simulations and machine learning algorithms. With the protocol suggested by Starr et al. (*23*) and the algorithms presented here, we foresee a completely computerized tool for suggesting improved antibody designs, with accuracy approximating DMS.

**Figure 2:**
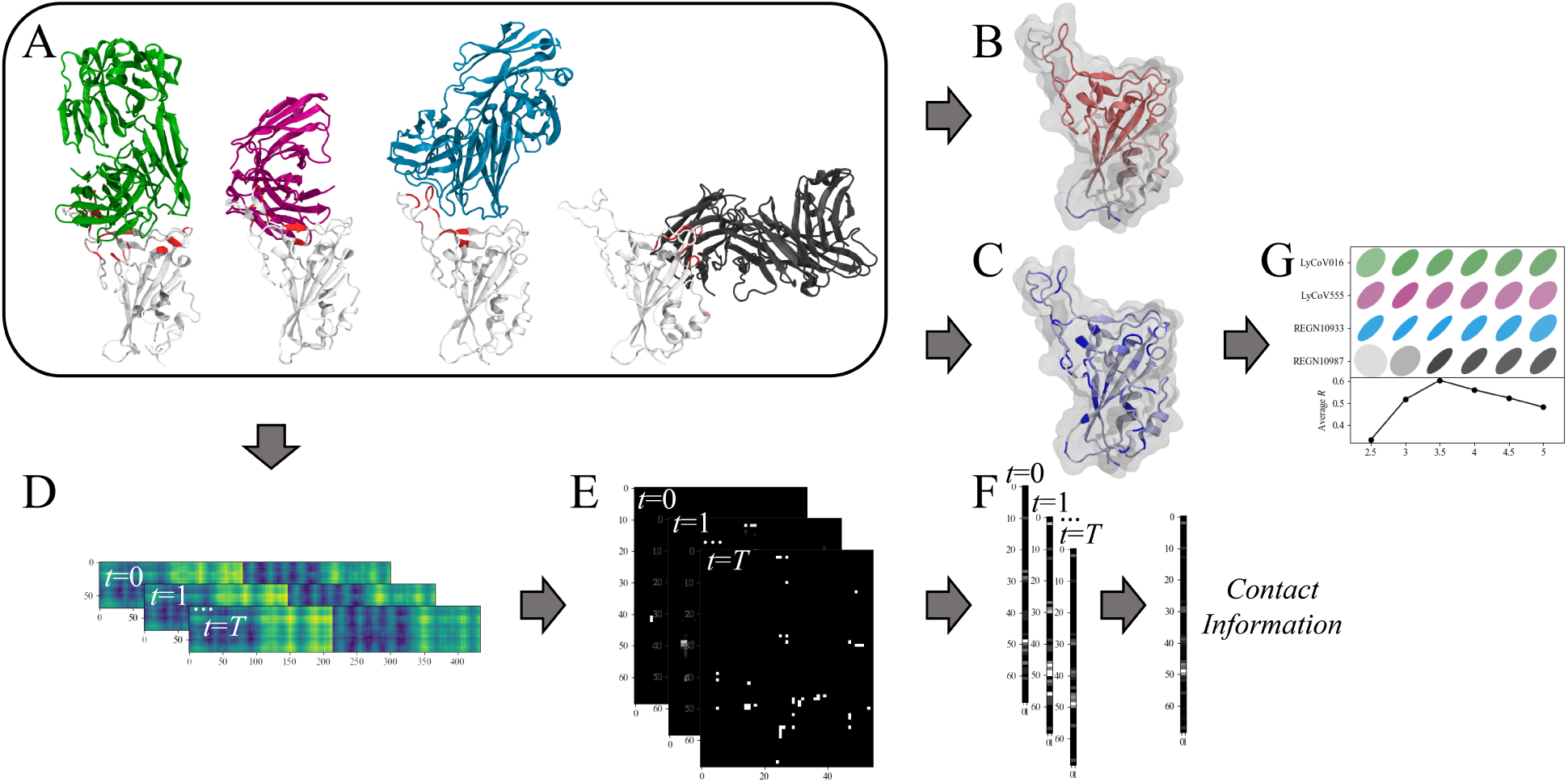
Schematics of how contact percentage and total contacts are calculated from structures. (A) Protein structures, where RBD is color-mapped with DMS total viral escape measurements (white being 0 and red being 1, normalized) (*21,22*). (B) RBD is colored with RMSF from one sample simulation, with white residues being more stable and red residues being more flexible. (C) RBD is colored with Wimley-White hydrophobicity, with white being more hydrophobic and blue being more hydrophilic. (D) Contact matrices are calculated from either crystal structures or simulations. (E) We convert contact matrices to binary contact matrix with a cutoff *c*. (F) We accumulated binary contact matrix in two ways to calculate contact information (contact percentage or total contacts). (G) We calculate the correlations between contact information and DMS total viral escape.

## Results and discussions

### Crystal structure contact information displays high correlation with DMS viral escape measurements

In the recent publication by Starr *et al*. (*23*), it was shown that crystal contacts may indicate mutational hotspots of the SARS-CoV-2 virus that could escape antibody treatment. Inspired by this observation, we systematically calculated the crystal contacts in two ways before calculating the correlation between crystal contacts and DMS total viral escape. The most straight-forward way of calculating crystal contacts is to directly use residue-residue distances between RBD and antibodies. By taking the minimal closest-heavy (see methods for definition) residue-residue distance for each of the RBD residues, we obtain an array of raw crystal contact distances (with a dimension of *I* × 1, where *I* is the number of RBD residues) that can be used to calculate the correlation with the DMS total viral escape (see Methods for the definition). In Figure S1, we compare the raw crystal contact distances extracted from the crystal structures (PDB ID: 7C01, 7KMG and 6XDG) and the DMS total viral escape measurement (*21,22*). Pearson correlation coefficients (*R* thereafter) are plotted with ellipse plots, where *R* = ±1 yields a diagonal line and *R* = 0 yields a perfect circle. *R* averaged over the four antibodies is determined to be -0.48, indicating a moderate negative correlation between the raw crystal contact distances and the DMS total viral escape. The negative correlation does not come as a surprise, since longer distances normally mean lower contributions from the residues to residue-residue interactions (*35*). The moderate correlation, however, is encouraging when linking structural properties to DMS total viral escape.

In an attempt to improve the aforementioned correlation, we explored another way of calculating crystal contacts. Instead of directly using distances, we set a cutoff (*c*) below which the residue-residue contact is *True*, above which the residue-residue contact is *False*. The resulting binary contact matrix is then condensed to the dimension of *I* × 1 by simply taking logical operator “OR” for each of the rows. Each element in this *I* × 1 contact array represents if the corresponding residue in RBD is in contact with any residues of the corresponding antibody in the crystal structure within a given cutoff distance. We term this measurement “contact percentage” (even though with one structure, all measurements are either 100% or 0%). In Figure 3A, we plotted *R* values with 6 different cutoff distances ranged between 2.5 Å and 5 Å. The average *R* peaks around 0.60 with a 3.5 Å as the cutoff (*c*), which is higher than *R* = -0.48 in absolute value calculated with raw residue-residue distances. Interestingly, in the literature, the closest-heavy residue-residue distances often have an empirical cutoff of around 4 Å with purposes such as residue contact likelihood prediction (*36*), protein folding recognition (*37*), coarse-grained interaction distance of MARTINI model (*38*), and analytical results 2*r*_0_, where *r*_0_ is the van der Waals radii of carbon (*39*). Our finding agrees with choices previously made by other researchers, even though our 3.5 Å yields better *R* than 4 Å in this study. It is noteworthy that, despite the moderately high correlation between binary crystal contact and DMS total viral escape, it is impossible to quantitatively rank the contribution of residues to protein-protein interactions using binary information. This is simply because of the noncontinuous nature of the binary contact matrix if there is only one structure to work with. In the following sections, we explore two ways to resolve this issue: by either taking a summation of the binary contacts for each of the RBD residues or expanding the sampled structures with simulations.

**Figure 3:**
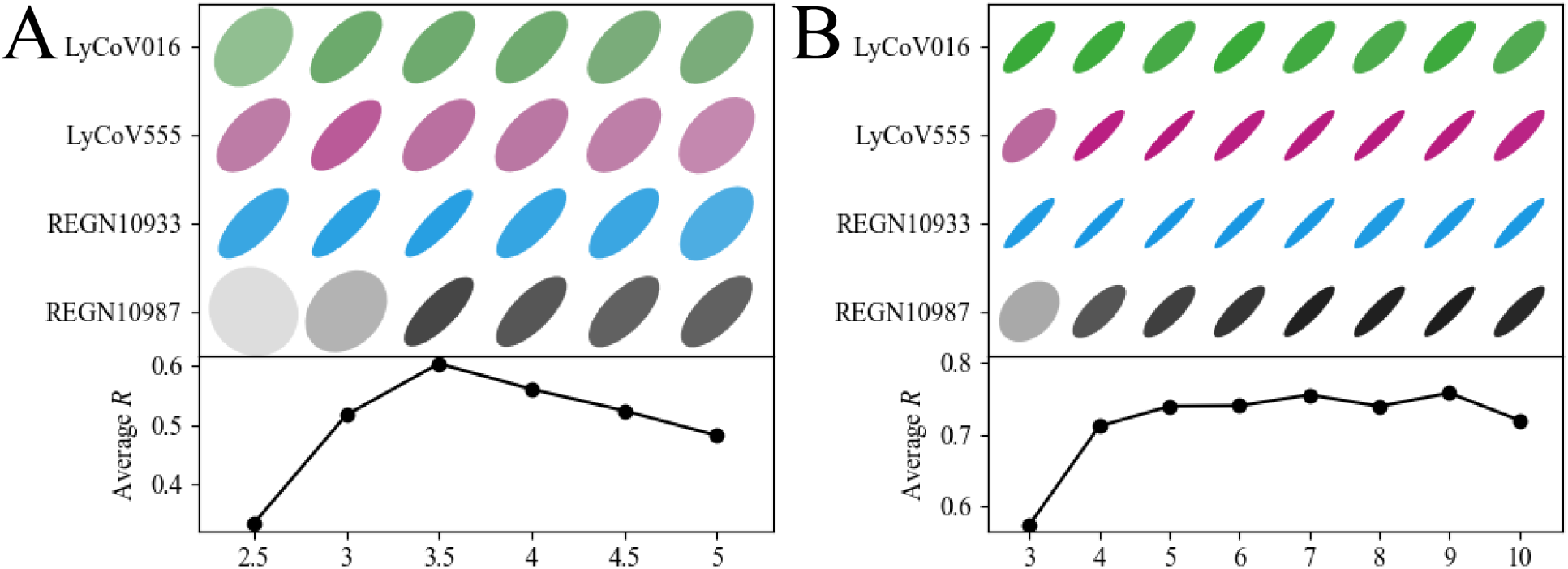
We calculated the average Pearson correlation *R* between contact information (using closest-heavy distances) from crystal structures and DMS total viral escape. Pearson correlation *R* values are plotted in ellipses (termed correlation ellipses) for each binding complex and expressed in two ways: ellipsoid aspect ratios and color saturation. Higher absolute values in *R* yield ellipses with higher aspect ratios and higher color saturation. Positive *R* values give diagonal ellipses while negative *R* values give anti-diagonal ellipses. To evaluate the choices of contact information, we calculated the averaged *R* values by averaging *R* values over all antibodies, plotted under the correlation ellipses. (A) Correlations between contact percentage and DMS total viral escape are calculated for crystal structures (PDB ID 7C01, 7KMG, 6XDG; note that REGN10933 and REGN10987 share the same PDB entry). Cutoff parameters *c* = 2.5 - 5 Å are compared (x axis). The correlation ellipses are always colored with green for LyCoV016, magenta for LyCoV555, blue for REGN10933, and gray for REGN10987. Accumulated *R* values over all antibodies are plotted under the correlation ellipses. (B) Correlations between total contacts and DMS total viral escape are calculated for crystal structures. Cutoff parameters *c* = 3 - 10 Å are compared (x axis).

To alleviate the difficulty of ranking residue contributions using binary information, we calculated the total number of residues in an antibody that are in contact with a residue in RBD by taking summations of the binary contacts, treating *True* as 1 and *False* as 0. We term this “total contacts”. In Figure 3B, we plot the *R* between total contacts and DMS total viral escape.

It is apparent that all *R* values calculated from total contacts are significantly higher than contact percentage. The highest *R* is 0.76, observed at both 7 Å and 9 Åm as the cutoff distances (*c*). With a standard deviation of the mean around 0.035, *R* values are statistically indistinguishable between 5 Å and 9 Å, where *R* ranges from 0.74 to 0.76. At *c* = 10 Å, *R* significantly drops to 0.72. It is interesting that the peak of *R* moves from *c* = 3.5 Å in Figure 3A to *c* = ∼7 Å in Figure 3B when the way we calculate the contacts is changed. More studies are required to explain this shift in details. Importantly, being able to use one static structure to achieve a strong correlation (with *R* = 0.76) against hundreds of experimental DMS measurements is exciting.

### Contact information calculated from simulations correlates well with DMS total viral escape

In order to assess if MD simulations can improve the aforementioned correlations, we carried out 3-4 replicas of simulations for all 4 antibody-RBD complexes, each lasting 400 ns. With a total 5.6 *µ*s of simulations, we calculated the binary contact matrix of each time frame the same way we calculated it with crystal structures. Similar to the aforementioned two approaches to condense binary contact matrix into contact arrays, we calculated the contact information (contact percentage and total contacts) for each of the time frames (Figure 2D, E, F). Finally, we took a simple average over all time frames to get the time-averaged contact information (Figure 2F).

The Pearson correlation *R*s are plotted in Figure 4 in ellipses. In Figure 4A, B, we compare the contact percentage with the total contacts, where all correlations are calculated with closest-heavy residue distances using all 400 ns of simulations. Similar to the case with crystal structures, *R* between simulated contact percentage and DMS total viral escape peaks around *c* = 3.6 Å at 0.66. This is higher than 0.60 calculated with crystal structures. Beyond *c* = 4.0 Å, *R* consistently drops. In regards to total contacts, *R* again peaks at both *c* = 7Å and 9Å with a value of 0.75, which is statistically identical to the values calculated from crystal structures (*R* = 0.76 at *c* = 7Å and 9Å). One significant difference is that instead of having a plateau from *c* = 5Å to 9Å for the calculated *R* using crystal structures, the plateau has been shrunk to *c* = 7Å to 9Å. Simulations seemingly do not affect the correlations as much as one would expect, which we will further investigate (see the next section where we find that short simulations improve the correlations). Overall, total contacts still yield significantly higher *R*s than contact percentage, indicating that total contacts are a better measurement than contact percentage for predicting DMS total viral escape.

**Figure 4:**
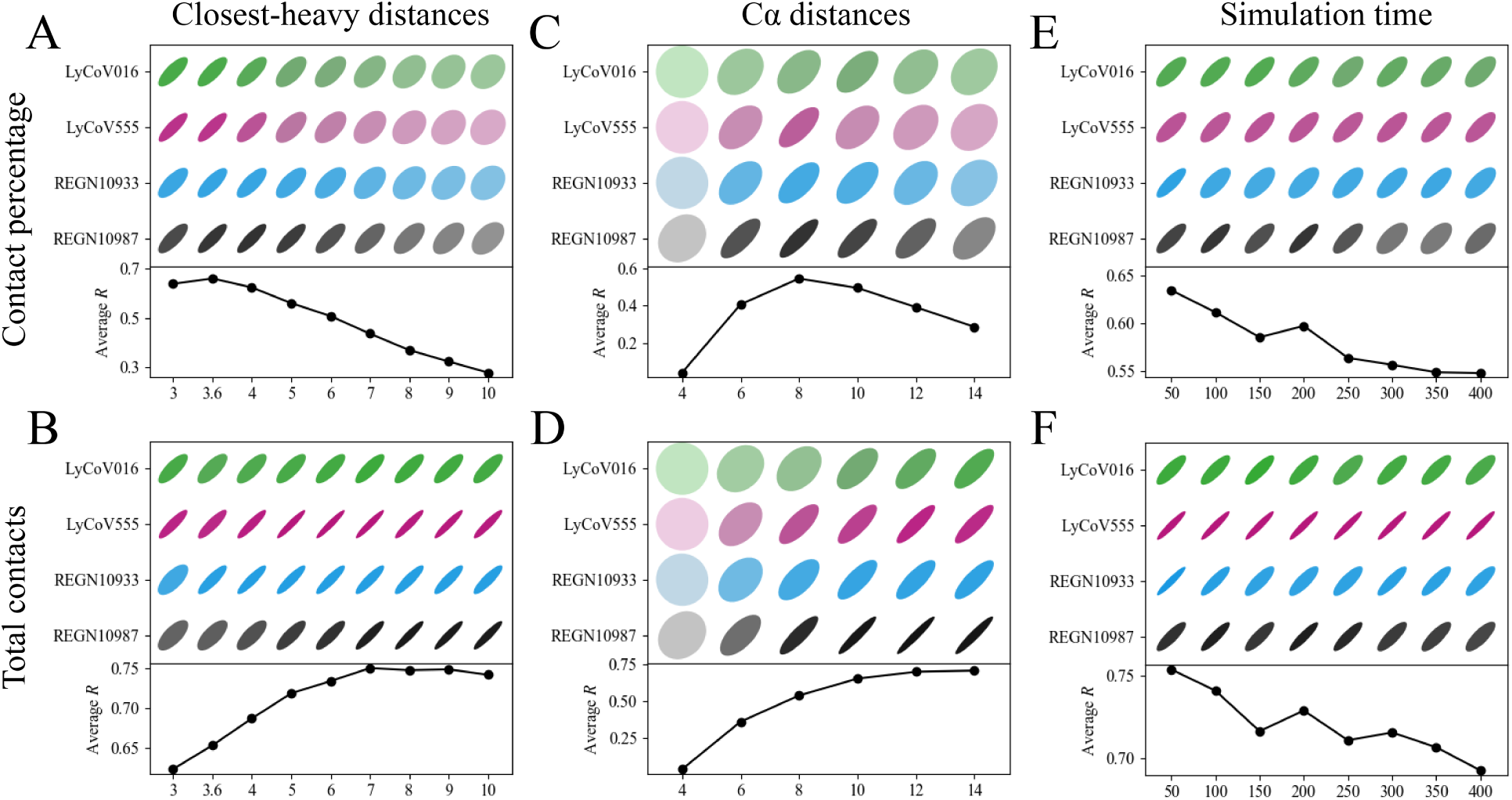
Similar to Figure 3, we calculated the Pearson correlation *R* between contact information and DMS total viral escape measurements and plotted them in correlation ellipses. But instead of using the static properties from the crystal structures, we averaged contact information over all time-frames from the simulations. In the first column (A, B), we use “closest-heavy distances” to determine whether two residues are in contact. We compare cutoff parameters *c* between 3 Å and 10 Å. In the second column (C, D), we use distances between alpha carbons to determine whether two residues are in contact. We compare cutoff parameters *c* between 4 Å and 14 Å. In the third column (E, F), we divide the simulations into chunks of 50 ns and compare the *R* values in these chunks, while using closest-heavy contacts and fixing *c* at 3.6 Å for contact percentage and 7 Å for total contacts, respectively. In the first row (A, C, E), we measure contact percentage, while in the second row (B, D, F), we measure total contacts.

In addition to the closest-heavy residue distances, we also explored the possibility of using just alpha carbons to calculate binary contact matrices. In Figure 4C, D, we plot the *R* values between contact information and DMS total viral escape using alpha carbon distances. As for contact percentage, *R* peaks at *c* = 8 Å with a value of 0.54, significantly lower than the 0.66 which we calculated with closest-heavy distances. For total contacts, *R* monotonically increases up to 14Å, where it mostly plateaus at 0.71, statistically lower than the 0.75 calculated with closest-heavy distances (*σ* = 0.4). It should not be surprising that peak *c* shifts to longer distances when contact matrices are calculated with alpha carbons, because residue radii are added in the process. The choice of 8 Å is well-known within the protein structure community when using alpha carbons as reviewed by Gromiha et al. (*35*) It is intriguing, however, that the shifted cutoff distances where *R* peaks are different for contact percentage (shifting ∼4.5 Å) versus total contacts (shifting ∼6 Å). If we consider the closest-heavy distances to be a “clean version” of residue-residue distances, alpha carbon distances are “blurry” because of the size differences of residues. Depending on the proteins, the average size of residues are slightly different, resulting in a minute shift in cutoff distances for the highest correlations in cases when we use contact percentage. From contact percentage to total contacts, we accumulate all binary contacts instead of maximizing the values. In this process, we add yet another layer of averaged residue distances, resulting in a more spread peak for *R*s, which might explain why the shifts are different for these two measurements. Because of the lower *R* values and the “blurry” nature of alpha carbon distances, in our machine learning models, we focus on closest-heavy distances (see below).

### Surprisingly, longer simulation time reduces the correlation with DMS measurements

To further analyze how MD simulations affect correlations, we divide the aforementioned 400 ns of simulations into 8 chunks, each lasting 50 ns. Using the same methods as before, we calculate and plot average *R* values for all complexes in Figure 4E, F. Note that cutoff *c* is selected at 3.6 Å for contact percentage and 7 Å for total contacts based on their peak values in Figure 4A, B. Despite two different ways of evaluating the correlations, *R* immediately drops after the first 50 ns of simulations, and continues to do so throughout the simulations. The biggest contributor to the drop is the complex of REGN10987-RBD, followed by REGN10933-RBD, with LyCoV555-RBD being the most consistent throughout the entire simulation. The average drop of *R* from the initial 0-50 ns to the final 350-400 ns is 0.09 (or 2*σ*) for contact percentage and 0.06 (or 1.3*σ*), which are both statistically significant. While more work is needed to further evaluate the effect of force field and simulation parameters on model predictability, it is important to note that longer simulations may not equate with better results.

Finally, we compared the *R*s calculated with total contacts in the first 25 ns of all simulations to the *R*s calculated with total contacts from the crystal structures (Figure S4). We obtained *R* = 0.78 at *c* = 9 Å from 25 ns of simulations (with *σ* = 0.04), versus *R* = 0.76 at *c* = 7 or 9Å from the crystal structures (with *σ* = 0.03). It appears that the highest average *R* calculated from the 25 ns simulations is not significantly higher than that from the crystal structures. In Figure S4, we plot the average *R*s averaged over all 25 ns simulations against cutoff *c* and compare them with *R*s averaged over all crystal structures against curoff *c*. We show that *R*s from 25 ns simulations are consistently higher than those calculated from crystal structures. This suggests that short simulations are able to improve the correlation between structural information and experimental measurement (DMS total viral escape).

### Simple machine learning models further improve the correlation between structural features and DMS total viral escape

We evaluated the possibility of further boosting the correlation between total contacts and DMS total viral escape measurements by using two regression models. In Figure 5, we plot the *R* values by using a linear regression (LR) model and a random forest regression (RFR) model. Both models are evaluated with five different input settings:

**Figure 5:**
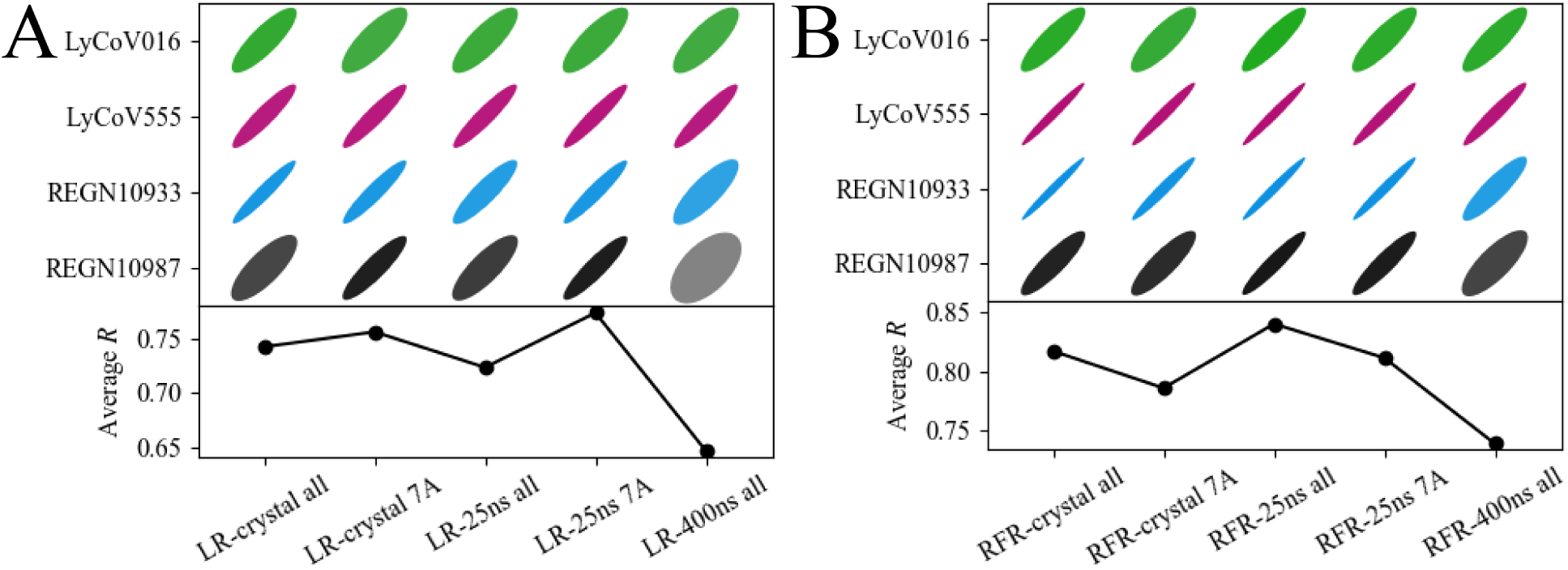
We used simple machine learning models to further boost the correlation between total contacts and DMS total viral escape measurements: (A) represents the linear regression (LR) model and (B) represents the random forest regression (RFR) model. We compared five methods of data preparation for both models. *R* calculated in the “Crystal all” columns accounts for total contacts of all cutoffs (*c* = 3, 4, 5, 6, 7, 8, 9, 10 Å) from crystal structures. *R* calculated in the “Crystal 7 Å “columns accounts for total contacts from crystal structures with only *c* = 7 Å. Similarly, *R* calculated in the “25ns all” columns and “25ns 7Å “columns accounts for total contacts measured from 25 ns of simulations. *R* calculated in the “400ns all” columns accounts for total contacts measured from 400 ns of simulations. Additional residue properties such as WW hydrophobicity and RMSF are also added for training, which are explained in detail in the Methods.

- “Crystal all” accounts for total contacts of all cutoffs (*c* = 3, 4, 5, 6, 7, 8, 9, 10 Å) from crystal structures and related Wimley-White hydrophobicity (WW) values.
- “Crystal 7 Å” accounts for total contacts from crystal structures with only *c* = 7 Å and related WW values.
- “25ns all” accounts for total contacts measured from 25 ns of simulations with all cutoffs and related WW and Root mean square fluctuation (RMSF) values.
- “25ns 7Å” accounts for total contacts measured from 25 ns of simulations with only *c* = 7 Å and related WW and RMSF values.
- “400ns all” accounts for total contacts measured from 400 ns of simulations with all cutoffs and related WW and RMSF values.

In Figure 5A, the LR model with crystal structures yields *R* values around 0.75, which is statistically the same as *R* = 0.76 using total contacts with cutoff *c* = 7 Å (Figure 4B). Comparatively, the LR model with 25 ns of simulations generates *R* = 0.77 (with only *c* = 7 Å), similar to *R* = 0.78 that was previously calculated with total contacts from 25 ns of simulations. The LR model with 400 ns of simulations is plotted as a baseline, reinforcing that shorter simulations seem to yield better correlations. In Figure 5B, RFR with crystal structures yields *R* values of 0.82 and 0.79 for “crystal all” and “crystal 7Å”, respectively, significantly higher than those obtained from total contacts (0.76 ± 0.03 with *c* = 7Å). RFR with 25 ns simulations yields *R* values of 0.84 and 0.81 for “25ns all” and “25ns 7Å”, respectively, also higher than those obtained from total contacts (0.77 ± 0.04 with *c* = 7Å). While the LR model shows no significant changes in correlation, the RFR model significantly improves correlation especially with total contacts of all cutoffs.

Focusing on the RFR results, it is notable that 25ns simulation *R* values, both in “all” and “7 Å” versions, are higher than the respective values achieved with crystal contacts. When considering all cutoffs the *R* value increases from 0.82 to 0.84; when considering only the cutoff *c* = 7 Å, we have an increase from 0.79 to the highest pick of 0.81. It is evident that both LR and RFR results confirm that the data derived from the shorter 25 ns simulations correlate better with DMS measurements than crystal structures.

## Conclusion

We present a simple measurement based on binary contacts between residues that correlates reasonably well with DMS viral escape measurements. With just crystal structures, we achieve an average *R* of 0.60 with contact percentage and 0.76 with total contacts. By equilibrating the crystal structures for 25 ns, we further improve the average *R* to 0.66 with contact percentage and 0.78 with total contacts. However, longer simulations seem to deteriorate the results, presenting a situation that calls for further study. By varying the parameters to obtain the binary contacts, we find that 3.6 Å and 8 Å are pivotal cutoffs to toggle the step functions when tallying the contacts for contact percentage and total contacts, respectively. These numbers match typical choices for calculating residue-residue contacts from previous studies (as discussed above). Finally, when we apply both the linear regression model and the random forest model, we further improve average *R* to 0.84. In conclusion, we find that by using a handful of structures, these easy-to-implement measurements correlate well with DMS viral escape measurements. We expect that with further improvement, this method will become a helpful tool in predicting viral mutants, designing antibodies, and solving other similar protein-protein interaction problems.

## Methods

### Molecular dynamics simulation

We use molecular dynamics simulations to collect dynamic contact information for four protein-protein interaction (PPI) complexes in this study. Within these PPI complexes, one binding partner is fixed to be the receptor binding domain (RBD) of SARS-CoV-2 spike glycoprotein (UniProtKB: P0DTC2). Four antibodies are examined as binding partners: LyCoV016 (*10*), LyCoV555 (*11*), REGN10933 (*12, 13*), REGN10987 (*12, 13*). The initial binding structures are taken from crystal structures deposited in RCSB Protein Data Bank (PDB): PDBID-7C01 (*10*) for LyCoV016, PDBID-7KMG (*11*) for LyCoV555, PDBID-6XDG (*12*) for REGN10933 and REGN10987, respectively. We perform standard structure cleaning procedures to the crystal structures, including adding missing residues based on structural similarities among these PDB files, adding missing atoms, fixing cis peptide bonds and fixing chirality with VMD 1.9.1 (*40*). Thereafter, we solvate the structures with TIP3P water and 0.15 M of NaCl using GROMACS2019 (*41*). All structures are prepared with the CHARMM36 force field (*42*). Before production runs, 10,000 steps of steepest descent energy minimization and 100 ps of equilibration MD simulations with canonical ensemble (NVT) and 1 fs time step are performed. Note that even though longer equilibrations are normally desired, we deliberately keep the equilibrations short to reveal a problem with the examined simulations (see discussion). Finally, 400 ns of production simulations are performed with 2 fs time step, 10 Å van der Waals switching distance, 12 Å smooth cutoff distance and 1 Å grid size for Particle Mesh Ewald calculations. All simulations are performed with GROMACS2019.

### DMS total viral escape

In the papers by Starr et al. (*21,22*), authors presented the percentage of virus escaping antibody treatment upon mutations on RBD. For each mutated RBD residue, 19 values were obtained for 19 possible mutations (sometimes missing 1 or 2). The total viral escape of residue *i* is a measurement that sums all 19 viral escape percentages for residue *i*.

### Structural measurements of PPI complexes

In the following sections, we present three structural measurements (*m*) that are calculated, compared, and utilized in this study. Those measurements are: Wimley-White hydrophobicity (WW), root mean square fluctuation (RMSF), and contact information calculated with residue-residue contact matrix (CM).

### Wimley-White hydrophobicity (WW)

Hydrophobicity is an important intrinsic property of an amino acid. We choose the Wimley-White scale of hydrophobicity (*43*) in this study as an independent measurement from distance measurements. Note that the viral escape measurement from DMS (*21, 22*) is a measurement of the percentage of viruses escaping antibody neutralization. Therefore, we convert the original Wimley-White scale in free energy to equilibrium constant. To capture the contribution of hydrophobicity to antibody escape, we make a list of hydrophobicity (denoted as WW) for any residue *i* in RBD by taking the natural logarithm of the original Wimley-White hydrophobicity in free energy (Δ*G*_Octanol*−*Interface_): WW_*i*_ = ln(ΔΔ*G*_*i*,Octanol*−*Interface_).

### Root mean square fluctuation (RMSF)

As a regular practice in a MD simulation, RMSF is often calculated to measure the flexibility of residues. First, we perform a structural alignment with backbone atoms of the part of the protein of interest (interfacial residues in RBD that are within 10 Å of the corresponding antibody) using VMD 1.9.1. Then, RMSF of residue *i* is simply calculated as 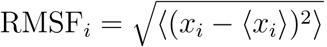, where *x*_*i*_ is the coordinates of the *α* carbon of residue *i*.

### Contact information calculated with residue-residue contact matrix (CM)

Residue-residue distances are calculated for each of the frames of the simulations for all PPI complexes in this study. The resulting *T* × *I* × *J* matrix from one simulation is denoted as the PPI contact matrix (CM), where *T* is the number of simulation frames, *I* is the number of residues in RBD, and *J* is the number of residues in an antibody. CM consists of *T* frames of submatrices CM_*t*_, where *t* ∈ [0, *T*]. Each CM_*t*_ consists of *I* × *J* elements, representing residue-residue distances. More specifically, CM_*t*_ = ((contacts_*r*1_),(contacts_*r*2_),…,(contacts_*rI*_)) where *ri* (*i* ∈ [1, *I*]) denotes the *i*th interfacial residue from RBD. (contacts_*ri*_) = (*x*1, *x*2, … *xJ*) where *xj* (*j* ∈ [1, *J*]) is the distance array between the *i*th interfacial residue and all residues of the examined antibody containing *J* residues. We then convert the distance contact matrices CM to binary contact matrices BCM by setting a cutoff value *c*. The elements in the matrices are converted to 0s and 1s depending on if the distances are above or below the cutoff *c*. In details, BCM_*t*_ = ((bincontacts_*r*1_),(bincontacts_*r*2_),…,(bincontacts_*rI*_)) and (bincontacts_*ri*_) = (*b*1, *b*2, … *bJ*) where *bj* = 1 if *xj* ≤ *c* and *bj* = 0 if *xj > c*. Finally, we remove all columns (not rows) in the binary contact matrices with only 0 values across all structures that are examined to save storage space and memory use.

To calculate the correlation between this structural measurement and DMS viral escape, the *I* × *J* dimension BCM_*t*_ needs to be collapsed to an one dimensional array of *I* measurements, denoted as contact arrays (CA_*t*_ = (*m*1, *m*2, …*mI*)). We experiment with two ways of doing so. The first way is to calculate whether a residue *i* in RBD is in contact with any residues in the antibody, formulated as *mi* = max((bincontacts_*ri*_)) where *i* ∈ (1, *N*). Then all BCM_*t*_ are averaged over all time frames, resulting in the percentage of time when a residue *i* in RBD is in contact with any residues in the antibody. We call this measurement “contact percentage”. The second way is to calculate how many residues in the antibody are in contact with a residue in RBD on average. To get this value, we calculate the total number of contacts at time frame *t* by using *mi* = sum((bincontacts_*ri*_)) and then average this value over all time frames. We call this measurement “total contacts”. “Contact information” in the main text refers to both “contact percentage” and “total contacts”, in order to make statements simpler.

Finally, to calculate the residue-residue distances, we experiment with an additional two methods. The first method is to use the so-called “closest-heavy” distance between two residues, which refers to the minimal distance of all heavy atom combinations (i.e. all atom combinations excluding hydrogen atoms) between the two residues: *x*_*i,j*_ = min(*d*_*C*(*i,j*),*heavy*_) where *x*_*i,j*_ is an element in CM_*t*_, *d* is a distance measurement, and *C*(*i, j*), *heavy* denotes all combinations of heavy atoms between the *i*th residue in RBD and *j*th residue in the antibody. The second method is to simply measure distances between alpha (*α*) carbons: *x*_*i,j*_ = *d*_*αi,αj*_. We compare these two methods of distance measurements in Figure 4.

Note that when calculating the CM of crystal structures, time dimension *T* = 1. Therefore, all measurements are based on one single structure if we only have crystal structures (Figure 3).

### Machine learning models

We tested simple Machine Learning models to predict the DMS total viral escape. In experiments with crystal structures we used WW hydrophobicity and the total contacts of all 8 cutoffs (*c* = 3, 4, 5, 6, 7, 8, 9, 10 Å) as input features of the models, thus the model has 9 input features. In experiments with MD simulations, we added the residue RMSF as an additional feature. Note that RMSF is generated by dynamics, which does not exist for crystal structures. We evaluated the machine learning models in a cross-protein fashion by blind tests, where, in turn, the data from one target protein-protein interactions is used as the test set and those related to the other three proteins are used to train the models. In our experiments, we tested two well known regression models using their scikit-learn (*44*) implementation:

- Linear regression (LR): Predictions are made by using a linear function *p*_*i*_ = *a*_0_ + *a*_1_ ∗ *m*_1_ + *a*_2_ ∗ *m*_2_ + …*a*_*n*_ ∗ *m*_*N*_ where *m*_*n*_ is the *n*th feature.
- Random Forest Regression (RFR): a popular supervised learning algorithm based on decision trees and bagging ensemble learning methods (*45*). We tuned the hyper-parameters (*max depth, max features, min samples leaf, min samples split* and *n estimators*) of the Random Forest model with cross-validation over the training set. For each experiment setting, we selected the combination of parameters that held better general results for all the four target protein-protein interactions.

## Author Contributions

L.Z. conceived and designed the study. L.Z. and C.Y. performed molecular dynamics simulations. L.Z. and G.D. collected and analyzed data. G.D. applied machine learning models. L.Z., G.D., and C.Y. interpreted results and co-wrote the manuscript. All authors contributed to the general discussion of the project and manuscript.

## Acknowledgement

The authors wish to acknowledge helpful discussions with Binquan Luan, Tien Huynh, Ruhong Zhou, and Guojing Cong. L.Z. gratefully acknowledges the financial support granted by the IBM exploratory life sciences council.

## Supporting information

### Residue hydrophobicity and simulation RMSF correlate poorly with DMS total viral escape

Hydrophobicity of residues was used often in force field developments and featurization of proteins. (*46, 47*) Here we use Wimley-White hydrophobicity (*43*) as an independent metric to assess its correlation with DMS total viral escape. Plotted in Figure S1, the *R* values are highly dependent on the protein-protein complex. While for LyCoV016, LyCoV555, and REGN10933 the correlation is almost non-existent, the correlation is moderately high for REGN10987 with *R* = 0.55. Considering the non-trivial correlation between hydrophobicity and DMS total viral escape, we included it as an input feature of our machine learning algorithm (see below).

A frequent practice in analyzing MD simulations is to calculate the root mean square fluctuations (RMSF) of the alpha carbons in simulated proteins. In this study, we calculate the RMSF of all residues in the RBD while aligning the binding complex. All *R* values between DMS total viral escape and RMSF have been plotted in Figure S1. The highest *R* based on absolute values calculated here is -0.30 which occurs in simulations with antibody LyCoV555. The average absolute value of *R* is 0.20, which suggests a weak correlation between RMSF and the experimental measurement. However, the average value of *R* is -0.08, indicating that signs of the correlation coefficients from individual simulations are random. Overall, the correlation between simulated RMSF and DMS total viral escape seems case-dependent. Therefore, based on our calculations, we strongly discourage the use of RMSF as a single metric to predict residues’ contributions to protein-protein binding.

As we have previously shown in a protein-ligand binding study (*31*) and a protein-peptide binding study (*32*), derived contact matrix measurements are uncorrelated with root mean square fluctuation (RMSF) measurements of the residues. Similar to the above, we calculate the average *absolute value* of *R* = 0.20 between contact percentages (or total contacts) and RMSF, which suggests a weak correlation. However, the average *value* of *R* is 0.01, again indicating that signs of the correlation coefficients from individual simulations are random. Therefore, we are confident that the derived contact information from our method is independent from RMSF calculations.

### RMSDs of simulations do not correlate with *R*

To examine how stable each of the binding complexes is when starting from the corresponding crystal structure, we perform root mean square displacement (RMSD) measurements of all backbone atoms (C, O, CA, and N) with crystal structures as references. The RMSDs are plotted against simulated time in Figure S2. Despite one rare binding conformation change for LyCoV016-RBD (after which the structure returns to normal), this complex mainly remains stable throughout all simulations with RMSD averaged at 0.25 nm. LyCoV555-RBD remains stable throughout all simulations as well, with RMSD averaged at 0.18 nm. Meanwhile, REGN10933-RBD and REGN10987-RBD deviate a bit more from the crystal structures, with RMSD averaged at 0.50 nm and 0.74 nm, respectively.

In the main text, we have discussed that with the goal of optimizing *R* between total contacts and DMS total viral escape, short simulations (25 ns) outperform static crystal structures, while longer simulations weaken the correlation. A natural question is whether the structural deviation (RMSD from crystal structures) directly affects *R* values. To answer this question, we plot RMSD against *R* by separating the 400 ns simulations into 8 chunks of 50 ns time-slices in Figure S3A. In Figure S3B, we plot RMSD against *R* only for the first 25 ns of the simulations. As indicated by Figure S3A, the correlation between RMSD and *R* is weakly negative (with a coefficient of -0.24). This may indicate that higher deviations possibly yield worse correlation (*R*) between total contacts and the DMS total viral escape, agreeing with common sense expectations. However, this weak correlation does not hold for 25 ns simulations, as presented in Figure S3B. Furthermore, with longer simulated time (data points with lower color saturation in Figure S3A), *R* decreases disregarding whether RMSD remains the same or slightly increases. Therefore, we conclude from the four examined binding complexes that the deviations from crystal structures do not necessarily indicate worse correlations between total contacts and DMS total viral escape. It will of course require further studies to validate this conclusion.

---

**Figure S1:**
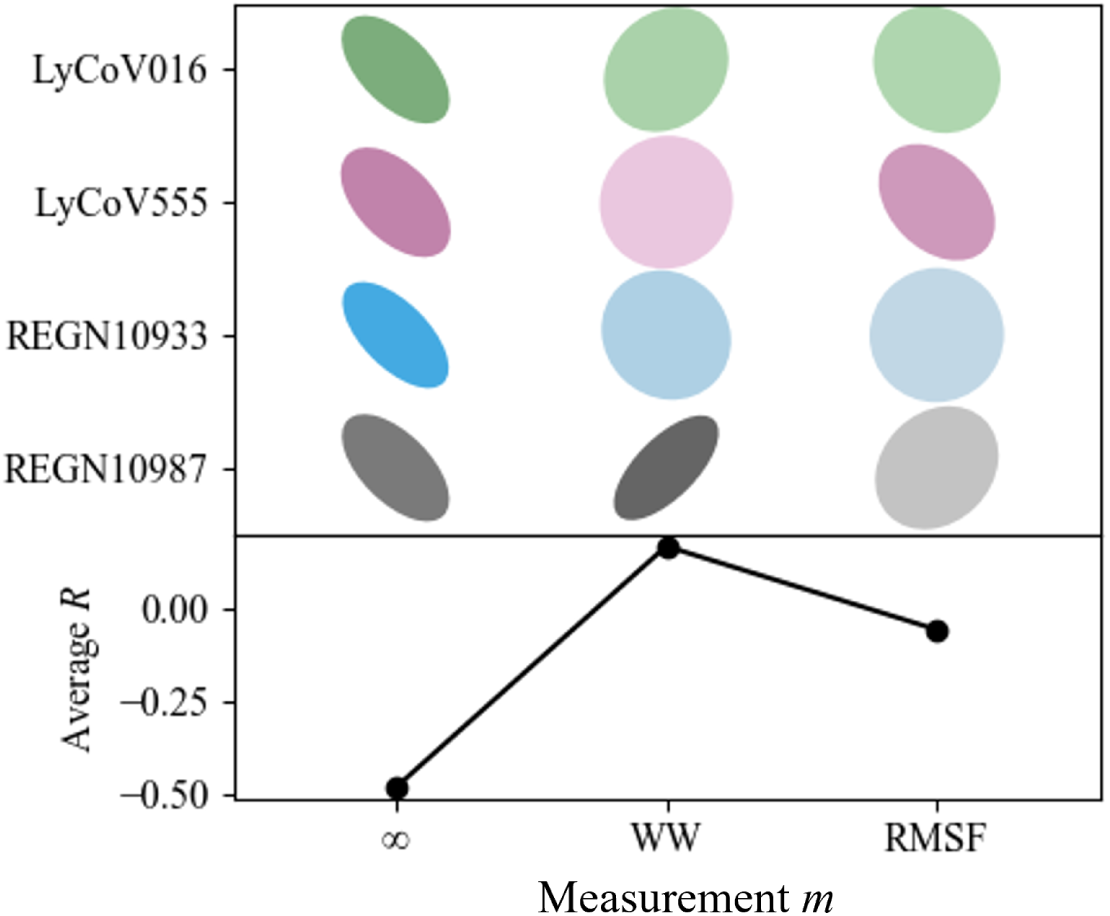
Correlation ellipses of 3 measurements: raw crystal contact distances (∞), Wimley-White hydrophobicity of residues (WW), and residue root mean square displacement of simulations (RMSF).

**Figure S2:**
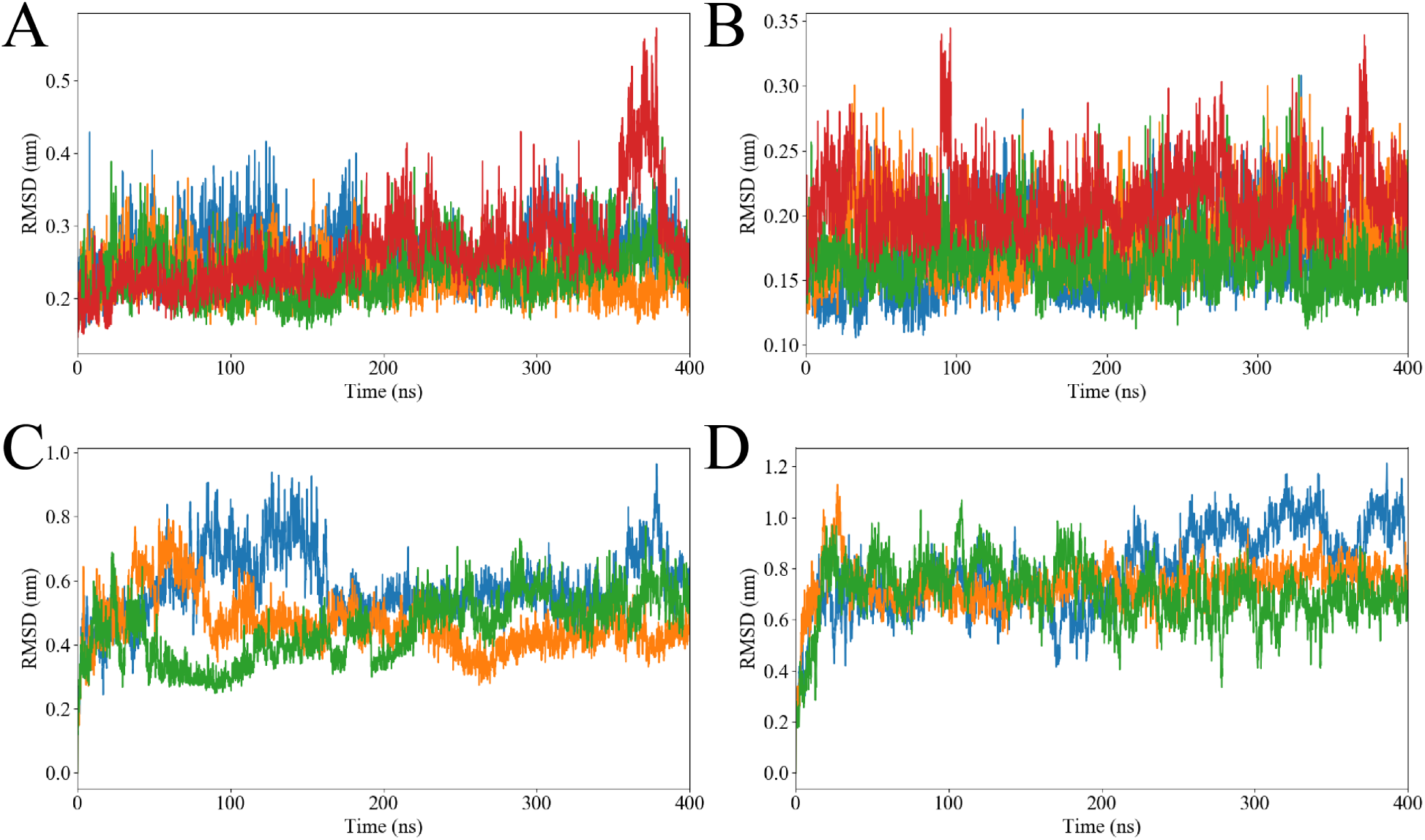
Root mean square displacement (RMSD) of the protein complexes for all simulations with LyCoV016-RBD in (A), LyCoV555-RBD in (B), REGN10933-RBD in (C), and REGN10987-RBD in (D).

**Figure S3:**
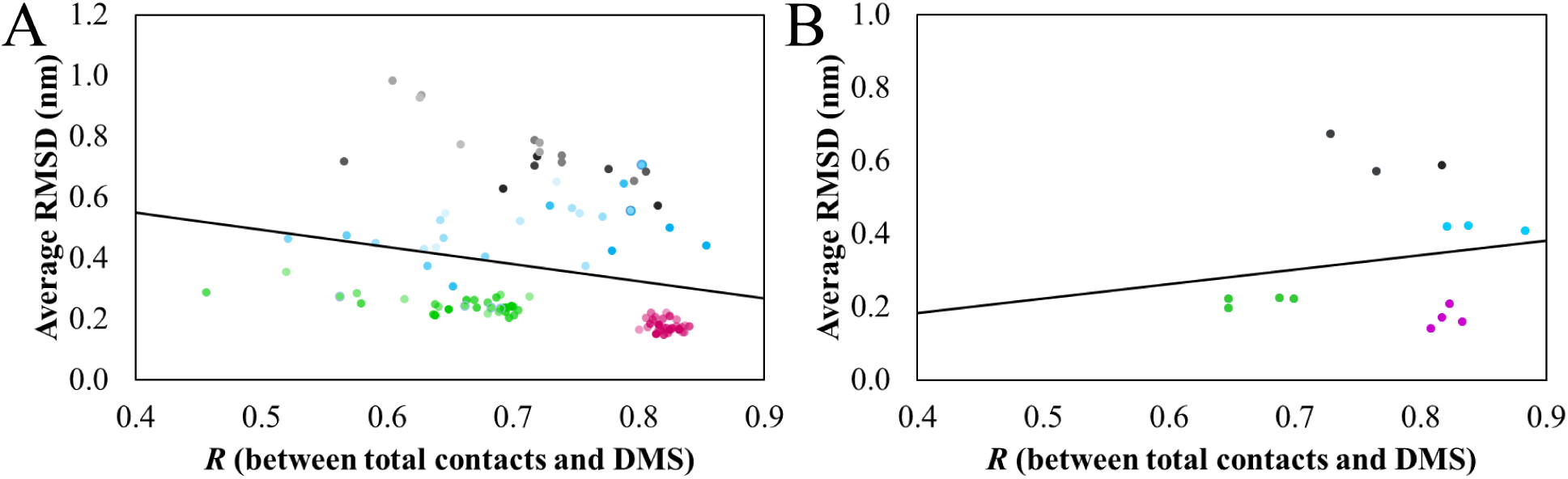
To assess how the stability of simulation affects *R* between total contacts and DMS total viral escape measurements, we calculated the correlation between RMSD (y axes) and *R* (x axes). In (A) we present the data collected over all 50 ns time-slices from all simulations. In (B) we plot the data points only from the initial 25 ns of all simulations. In all plots, data points are colored green for LyCoV016-RBD, magenta for LyCoV555-RBD, blue for REGN10933-RBD, and black for REGN10987-RBD. Saturation of the data points indicate how much simulated time has elapsed, with 100% corresponding to 0-50 ns and 40% corresponding to 350-400 ns.

**Figure S4:**
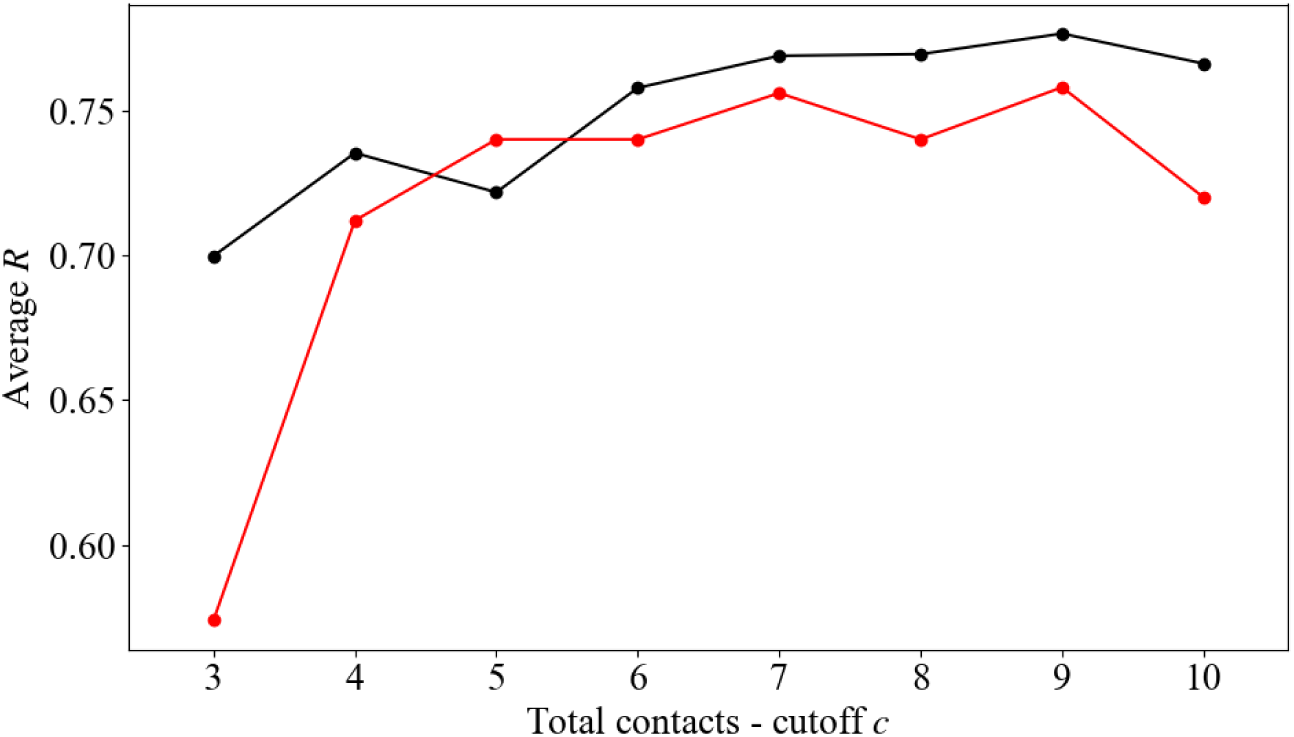
Average correlation coefficients *R* (calculated between total contacts and DMS total viral escape measurements and averaged over four examined binding complexes) are compared between crystal total contacts (red) and simulated total contacts, with only initial 25 ns of simulations (black) at all cutoffs (*c* = 3 - 10 Å).

